# Young children combine sensory cues with learned information in a statistically efficient manner: But task complexity matters

**DOI:** 10.1101/776260

**Authors:** Vikranth R. Bejjanki, Emily R. Randrup, Richard N. Aslin

## Abstract

Human adults are adept at mitigating the influence of sensory uncertainty on task performance by integrating sensory cues with learned prior information, in a Bayes-optimal fashion. Previous research has shown that young children and infants are sensitive to environmental regularities, and that the ability to learn and use such regularities is involved in the development of several cognitive abilities. However, it has also been reported that children younger than 8 do not combine simultaneously available sensory cues in a Bayes-optimal fashion. Thus, it remains unclear whether, and by what age, children can combine sensory cues with learned regularities in an adult manner. Here, we examine the performance of 6-7-year old children when tasked with localizing a ‘hidden’ target by combining uncertain sensory information with prior information learned over repeated exposure to the task. We demonstrate that 6-7-year olds learn task-relevant statistics at a rate on-par with adults, and like adults, are capable of integrating learned regularities with sensory information in a statistically efficient manner. We also show that variables such as task complexity can influence young children’s behavior to a greater extent than that of adults, leading their behavior to look sub-optimal. Our findings have important implications for how we should interpret failures in young children’s ability to carry out sophisticated computations. These ‘failures’ need not be attributed to deficits in the fundamental computational capacity available to children early in development, but rather to ancillary immaturities in general cognitive abilities that mask the operation of these computations in specific situations.

**Research Highlights:**

- Young children are sensitive to, and can learn, environmental regularities. Can they also utilize such learned regularities in a statistically efficient manner?
- We demonstrate that 6-7-year olds are capable of learning and utilizing regularities in a statistically efficient fashion, and in a manner indistinguishable from adult behavior.
- However, variables such as task complexity can influence young children’s behavior to a greater extent than that of adults, leading their behavior to look sub-optimal.
- These findings have important implications for how we should interpret failures in young children’s ability to carry out sophisticated computations.

## Introduction

Human observers operate in a world of sensory uncertainty, created by noise or processing inefficiencies within each sensory modality, or by variability in the environment (Knill & Pouget, 2004). Given the presence of such internal and external uncertainty, task performance is necessarily limited by the quality of the sensory information available on each trial. However, if the observer has access to multiple simultaneous sources of information (or cues) about the stimulus of interest (within or across sensory modalities), combining the information across cues in a statistically optimal fashion can lead to reduced uncertainty and improved performance (Bernardo & Smith, 2009; Cox, 1946; Jacobs, 2002; Knill & Pouget, 2004; Yuille & Bulthoff, 1996). Indeed, human adults are adept at mitigating the influence of sensory uncertainty by integrating multiple sources of information in an approximately Bayes-optimal fashion (i.e., by weighting each cue by its relative reliability). Numerous studies have shown that adults combine multiple cues within and across sensory modalities in such a Bayes-optimal fashion (Alais & Burr, 2004; Bankieris, Bejjanki, & Aslin, 2017; Ernst & Banks, 2002; Hillis, Ernst, Banks, & Landy, 2002; Jacobs, 1999; Knill & Saunders, 2003; van Beers, Sittig, & Gon, 1999). Of course, this strategy is only tenable when the observer has access to multiple simultaneous cues about the stimulus of interest. In situations in which the observer does *not* have access to multiple simultaneous cues, the influence of sensory uncertainty can nevertheless be mitigated by learning the underlying statistical properties (i.e., the generative model) for the stimulus of interest, and efficiently combining this learned prior knowledge with the sensory information available on each trial. Indeed, a large body of evidence suggests that human adults can improve their performance by combining uncertain sensory information with learned prior information in an approximately Bayes-optimal manner (i.e., by weighting each source of information by its relative reliability; (Bejjanki, Knill, & Aslin, 2016; Berniker, Voss, & Kording, 2010; Hudson, Maloney, & Landy, 2007; Jazayeri & Shadlen, 2010; Kording & Wolpert, 2004; Kwon & Knill, 2013; Stocker & Simoncelli, 2006; Tassinari, Hudson, & Landy, 2006; Vilares, Howard, Fernandes, Gottfried, & Kording, 2012).

Despite the preponderance of evidence supporting human adults’ ability to carry out such sophisticated computations, relatively less is known about the developmental time-course of this ability. Recently, some studies have reported that young children differ substantially from adults in their ability to mitigate the influence of sensory uncertainty by utilizing multiple sources of information (Barutchu, Crewther, & Crewther, 2008; Chambers, Sokhey, Gaebler-Spira, & Kording, 2018; Gori, Del Viva, Sandini, & Burr, 2008; Nardini, Bedford, & Mareschal, 2010; Nardini, Jones, Bedford, & Braddick, 2008; Petrini, Remark, Smith, & Nardini, 2014). For instance, several studies have shown that when presented with multiple simultaneous cues children younger than 8-12 years of age combine these cues in a sub-optimal manner – their behavior is often dominated by one of the cues irrespective of its relative reliability. The source of this dramatic discrepancy between the behavior of young children and adults remains unclear. What is the maturational trigger, or the threshold of accumulated environmental input, that enables 8- to 12-year olds (but not younger children) to demonstrate adult-like behavior? Some have argued that the basis for this sub-optimality is that sensory systems are rapidly changing in children younger than 8, which results in children prioritizing the use of one cue (or a subset of cues) to calibrate others (i.e., cue recalibration), thus precluding optimal cue combination (Gori et al., 2008). Others have suggested that the underlying computational process may be different during early development, which would manifest in optimal weights that are not solely dependent on cue reliability. For instance, examining the integration of stereo and disparity cues to surface slant, Nardini et al. (2010) found that children younger than 8 seem to be able to keep the two cues separate (as opposed to adults who experienced mandatory fusion of the two cues), thereby allowing them to prioritize the fastest-available cue, and to better detect conflicts between rapidly changing sensory modalities.

Notably, most of these prior studies have used tasks in which young children are presented with multiple simultaneously available cues (within or across sensory modalities). Little previous work has been devoted to the related question of whether young children’s behavior is similarly sub-optimal when tasked with integrating uncertain sensory information with prior information (but see Chambers et al. (2018)). This is an important gap in the literature for several reasons. First, as described above, adults have been shown to use similar mechanisms both when integrating simultaneously available cues, and when integrating prior information with uncertain sensory information. In both cases, adults’ behavior is consistent with the predictions of a Bayes-optimal observer that weights each available cue by its relative reliability. Thus, it is important to know whether children, like adults, utilize similar computational mechanisms (i.e., weight each available cue by its relative reliability) both when integrating simultaneously available cues, and when integrating learned prior information with uncertain sensory information. Second, there is substantial evidence that young children, and even infants, are sensitive to the statistical properties of stimuli in their environment (Fiser & Aslin, 2002; Gopnik, Sobel, Schulz, & Glymour, 2001; Jusczyk & Aslin, 1995; Kirkham, Slemmer, & Johnson, 2002; Kuhl & Meltzoff, 1982; Neil, Chee-Ruiter, Scheier, Lewkowicz, & Shimojo, 2006; Saffran, Aslin, & Newport, 1996; Smith & Yu, 2008; Xu & Garcia, 2008), and the ability to appropriately learn and use such environmental regularities has been shown to be involved in the development of several cognitive abilities (e.g., object recognition, language acquisition, causal inference etc.). Furthermore, a large body of evidence has supported the notion that young children and infants are able to utilize sophisticated computational abilities to deal with uncertainty in their environments (Bonawitz, Denison, Griffiths, & Gopnik, 2014; Saxe, Tenenbaum, & Carey, 2005; Schulz, Goodman, Tenenbaum, & Jenkins, 2008; Xu & Garcia, 2008). Thus, a failure to combine information efficiently cannot be the result of an insensitivity to that information per se. Third, mechanisms implicated in the sub-optimal behavior observed when integrating multiple simultaneously available cues (e.g., cue recalibration or prioritizing the earliest available cue) are less likely to be impediments to efficiently integrating sensory and prior information because there are no obvious cue-conflicts. Indeed, the presence of cue-conflicts would lead to further difficulties because they require the observer to infer the extent to which both cues pertain to the same underlying stimulus (or source) – only cues pertaining to the same source should be integrated. This “causal inference” problem (Kayser & Shams, 2015; Körding et al., 2007; Rohe & Noppeney, 2015; Shams & Beierholm, 2010) is much less relevant in situations involving the integration of prior and likelihood information.

Here, we examine the computational mechanisms used by 6-7-year old children when tasked with localizing a ‘hidden’ target by combining uncertain sensory information with prior information that could be learned over repeated exposure to the task. We provide evidence that in such situations, children younger than 8 years of age are capable of demonstrating behavior that is consistent with the predictions of a Bayes-optimal observer, and indistinguishable from adult behavior. Importantly, we also show that variables such as the complexity of task-relevant statistical distributions can influence young children’s behavior to a greater extent than that of adults, potentially leading their behavior to look sub-optimal as they revert to simpler implicit strategies for guiding their performance in the face of uncertainty. Our findings build upon prior research showing that young children and infants are sensitive to environmental regularities, and can utilize sophisticated computational abilities to deal with uncertainty, to show that 6-7-year olds are also capable of utilizing learned regularities in a statistically efficient manner. This renders them capable of the “variance reduction” that is a signature outcome of Bayesian inference, when combining prior and likelihood information, albeit in situations involving reduced complexity. Our findings have important implications for how we should interpret failures in young children’s ability to carry out sophisticated computations. These ‘failures’ need not be attributed to deficits in the fundamental computational capacity available to children early in development, but rather to ancillary immaturities in general cognitive abilities that mask the operation of these computations in specific situations.

## Experiment 1

### Method

#### Participants

Eight 6- to 7-year old children (3 male; 5 female) participated in this experiment. Participants’ ages ranged from 7 years and 32 days to 7 years and 292 days (M = 7.25 years). One additional participant was eliminated from the study due to an inability to follow instructions. Participants provided informed assent, and their caregivers provided informed written consent. Participants were given small prizes in the form of stickers, small toys etc., during their participation, while caregivers were compensated $10 per hour, for their time. The University of Rochester’s institutional review board approved all experimental procedures.

#### Task description

Participants completed a spatial localization task that was identical to that used previously by Bejjanki et al. (2016) with adults (Experiment 2 in that study). On each trial, participants estimated the location of a ‘hidden’ target by touching the appropriate location on a touch-sensitive display (see Fig. 1). The horizontal and vertical co-ordinates of the target location on each trial were independently sampled from a location-contingent mixture distribution over two underlying isotropic 2-D Gaussian distributions. These underlying Gaussian distributions differed in their mean locations (one was centered in the left half of the display while the other was centered in the right half), and relative variances (one had an SD of 40 pixels while the other had an SD of 20 pixels). Across participants, the distributions were always centered at the same locations on the left and right of the display, but the variance assigned to each was counter-balanced (with the higher variance distribution centered in the left half of the display for 50% of the participants, and in the right half for the other 50%).

**Figure 1:**
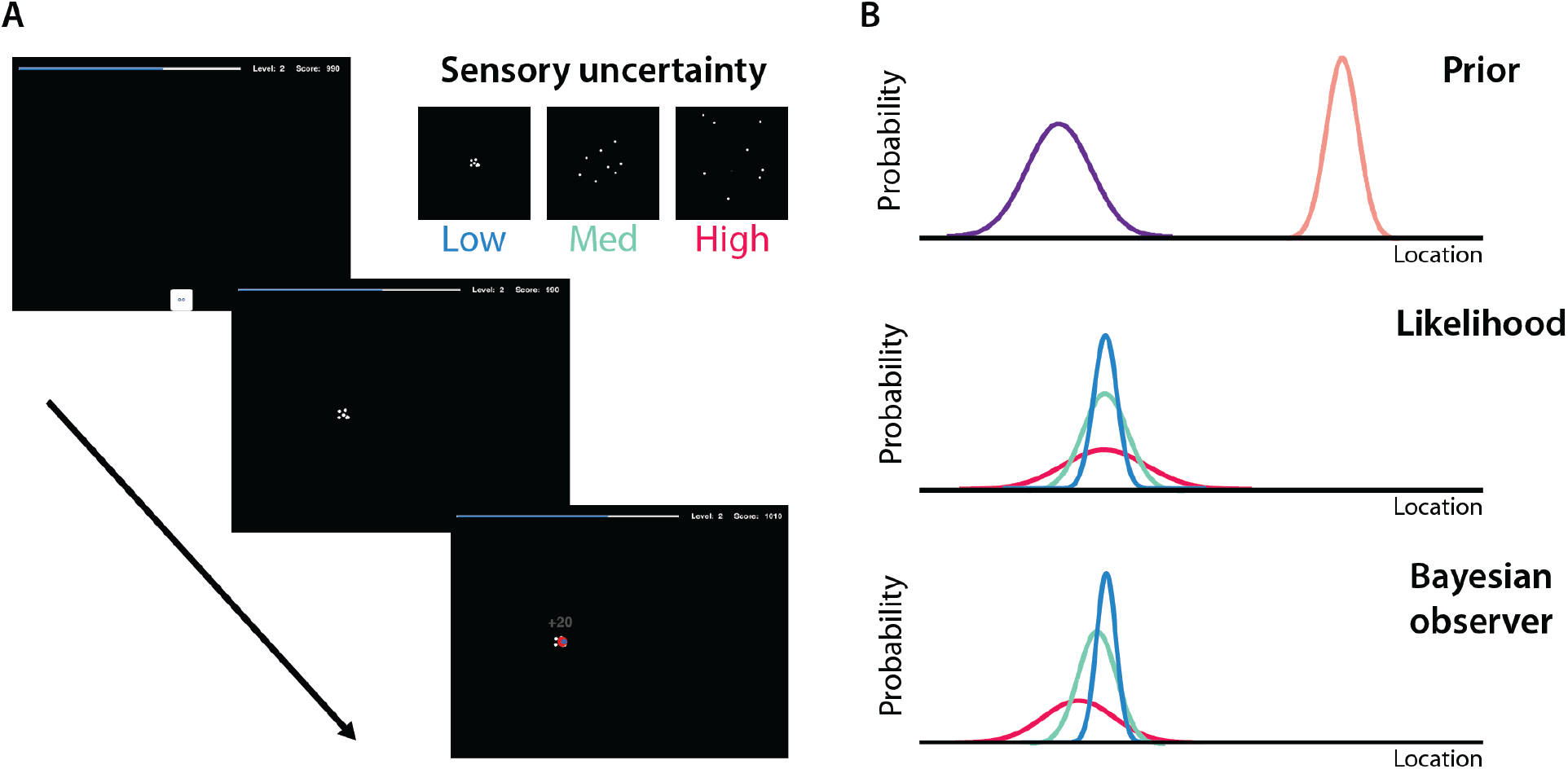
Learning and inference in a spatial localization task. (A) An illustration of a typical trial. Participants estimated the location of a ‘hidden’ target, randomly sampled on each trial from a mixture of two underlying Gaussian distributions. Upon touching a ‘GO’ button, they were presented with uncertain sensory information in the form of a dot cluster, centered on the target location, and subject to one of three levels of variability (low variability shown here; see inset for an illustration of the three levels). Feedback was provided post-touch. (B) An illustration of Bayes-optimal behavior. Considering the example of a ‘hidden’ target drawn from the more variable underlying distribution, the ideal observer would estimate the target location by learning the mean and variance of the prior (the task-relevant underlying distribution) and integrating this knowledge with the likelihood (the sensory information).

On each trial, participants were presented with uncertain sensory information about the target location in the form of a cloud of eight small white or green dots (depending on the underlying distribution that the target was drawn from). This cloud was independently generated on each trial by drawing samples from a separate 2-D isotropic Gaussian distribution centered at the true target location for that trial (i.e., the location of the target drawn from the underlying mixture model). The variance of this cloud of dots was manipulated to generate three levels of sensory uncertainty: the dots were drawn from a distribution that either had low variance (SD of 10 pixels), medium variance (SD of 60 pixels) or high variance (SD of 100 pixels; see inset of Fig. 1A). Thus, on each trial, participants had access to sensory information about the location of the hidden target (i.e., the cloud of 8 dots) as well as prior information about the likely locations of the hidden target, learned from feedback received in previous trials. Importantly, because the hidden target location was drawn from a Gaussian distribution (with SD of 20 or 40 pixels), the centroid of the cluster of dots was almost always discrepant from the average location of the hidden target. This spatial discrepancy allows us to estimate the weights attributed to the likelihood and prior information as a function of the reliability of each source of information.

Participants were also presented with a randomly interleaved ‘prior-only’ condition, in which no sensory information was presented (except for a briefly-flashed white or green box indicating from which of the two underlying distributions the target on that trial was drawn – see Fig. S1). In this condition, observers thus had to estimate the target location based solely on their learned knowledge (up to that point in the experiment) about where the target was likely to occur (i.e., based on their prior knowledge). After participants provided their response, feedback was provided via two new dots, one at the touched location and the other at the true target location. Participants also received feedback in the form of numerical points, with the point magnitude varying based on their accuracy.

#### Procedure

Before the start of the experiment, participants were provided with the following information:

You will be playing a game on an iPad. Imagine you are at a fair and you are playing a game in which there is an invisible bucket that you are trying to locate on the iPad screen. Sometimes the bucket is located on the left side of the screen and at other times it is located on the right side of the screen. Because the bucket is invisible, you can’t see where it is. However, each time you press “GO”, you’ll see some dots on the screen. You can think of these dots as guesses for where the bucket is. Now, you don’t know which (if any) of these dots actually show the location of the bucket. All of the dots could be just random guesses, which means they don’t actually tell you where the bucket is.

Sometimes you might be the first person to play the game, so you won’t see any dots on the screen when you press “GO”. Instead, you will briefly see a box on the screen, which will indicate the side of the screen that the invisible bucket is located at – the bucket will be somewhere within that box.

Your job each time is to try to guess where the bucket actually is, by touching the screen right where you think it is. When you touch the screen, you’ll see two more dots. One dot will be small and blue; this is where you guessed. One dot will be big and red; this is where the bucket actually was hiding. So, when you see the big red dot, that is where the invisible bucket actually was.

Then you’ll restart the game by pressing “GO” again, and try to find the new location of the bucket. The bucket will not always be in the same place, so pay attention to figure out where it might be. The closer you are to the bucket, the more points you get. If the blue dot and the big, red dot are right on top of each other, you get 20 points! If the blue and red dots are close to each other, you get 5 points – you just need to try a bit harder! If your guess, the blue dot, is very far away from the red dot, you get 0 points. Try to get as many points as you can, because you can go up in levels as you get more points, and win prizes. The closer your guesses are to the invisible bucket, the more points and prizes you get! Remember the bucket moves around; it is not always in the same place so you should try to figure out where it might be each time. Try your hardest to touch the right spot, so you get more points and prizes!

To ensure comprehension, this information was presented as a dialogue (rather than as a fixed script). During this dialogue, the experimenter frequently checked to ensure that the participant was following along, and participants were provided with ample opportunities to ask questions. Once participants confirmed that they understood the task, they were seated at a comfortable viewing distance from an Apple iPad (iPad 2; Apple Inc., Cupertino, CA), with a resolution of 768 (vertical) x 1024 (horizontal) pixels. Participants provided responses using their finger, having been instructed to pick a favorite finger and to consistently use that finger throughout the experiment. At the start of each trial, they were presented with a “GO” button centered at the bottom of the display. Once they touched this button, the trial started with the presentation of either a cluster of white or green dots, or a briefly flashed white or green box (on prior-only trials). Participants were told to estimate the location of the hidden target as rapidly and accurately as possible. Feedback was provided post-touch.

We implemented several measures to ensure that 6- to 7-year old children found this task engaging and fun. For instance, participants were able to earn points on each trial, with the magnitude of the points varying based on their accuracy. The game also included the opportunity to ‘level up’: when a participant accumulated 600 points on each level, they were shown a congratulatory screen and provided with small prizes (e.g. stickers). Progress towards the next level was always shown at the top of the screen, via a progress bar. This ability to ‘level up’ was used solely to motivate participants and did not represent any change in the experimental procedure. In addition, the experiment was designed to be self-paced such that each trial only began when the participant touched the “GO” button. Participants were encouraged throughout the study to take breaks as needed, and every time a participant leveled up, they were provided with the opportunity to take an extended break. During these breaks, participants were given the opportunity to briefly leave the room or engage in alternative tasks (such as coloring or playing with toys or stickers). Each participant completed 1600 trials across two one-hour sessions – each session involved 800 trials and was run on a different day.

#### Data Analysis

To quantify the computational mechanisms used by 6- to 7-year old children in this task, we estimated the extent to which their behavior depended on the sensory information available on each trial. Furthermore, we sought to characterize learning-induced changes in participants’ behavior, as a function of exposure to the task, by splitting the total number of trials into four temporal bins, each of which included 400 trials (with the eight conditions randomly interleaved in each bin).

For the trials in which sensory information was available, we used linear regression to compute the weight (*w*_*l*_) assigned to the centroid of the cluster of dots (the likelihood), with the weight assigned to the mean of the underlying target distribution for that trial (the prior) being defined as (1-*w*_*l*_). Thus, on each trial, given the centroid of the sensory information (*μ*_*l*_), the mean of the underlying target distribution (*μ*_*p*_) and participants’ estimate for the target location 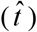, the weight assigned by participants to the centroid of the sensory information (*w*_*l*_) was estimated using:

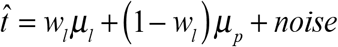

For the trials in which no sensory information was available (i.e., the ‘prior-only’ conditions), we computed participants’ mean responses across all trials in each temporal bin and for each prior condition. We focused on participants’ performance in the vertical dimension, to eliminate the potential influence of variability that may have been introduced by an interaction between participants’ handedness and the horizontal separation of prior locations. Moreover, participants’ performance in the horizontal dimension was similar to their performance in the vertical dimension – we found no interaction between the horizontal/vertical dimension and exposure, across prior and likelihood conditions (all *ps* > 0.05).

### Results

#### Learning and inference in a spatial localization task

On each trial of this task, participants had to estimate the location of a ‘hidden’ target based on two available sources of information: the sensory cue (the cluster of eight dots), and the history of feedback obtained from responses on all previous trials (the prior; see methods). The spatial distribution of each dot cluster (low, medium, or high variance) provided a more or less reliable estimate (reliability being inversely proportional to the cluster variance) of the true target location (Fig. 1A). Furthermore, dots in the cluster were either white or green, depending on the underlying distribution that the target was drawn from (narrow or broad), thereby providing a second cue (in addition to the location of the dot cluster) as to which of the two underlying distributions the target on that trial was drawn from. In order to independently estimate the extent to which participants learned the statistical properties of the underlying Gaussian distributions, we also randomly interspersed a ‘prior-only’ condition in which they localized the target in the absence of any sensory information (with the exception of a briefly-flashed white or green box indicating which of the two underlying distributions the target on that trial was drawn from – see Fig. S1). Participants thus had to estimate the location of the target based solely on their learned knowledge (up to that point in the experiment) about where the target was likely to occur, thereby allowing us to estimate their evolving knowledge of the location-contingent generative model, as a function of exposure to the task.

There are several computational models that participants could potentially use to carry out this task. The simplest strategy would be for participants to base their performance solely on the sensory information available on each trial, by choosing the centroid of the dot cluster as their estimate for the target location. This model (‘Model 1’) predicts a weight of 1 to the sensory information irrespective of the reliability of the sensory information or the reliability of the underlying target distribution for that trial. Furthermore, it predicts no change in participants’ weights as a function of exposure to the task, since the sensory information is independently drawn on each trial. This model is sub-optimal: as sensory uncertainty increases, the probability of the cluster centroid corresponding exactly to the true target location decreases; and by ignoring the underlying distributions, participants cannot exploit the fact that not all spatial locations are equally likely.

A slightly more sophisticated strategy involves participants taking the underlying distributions into account, but only learning and utilizing their mean locations while ignoring their relative variances (i.e., relative reliability) – learning the means is statistically easier and requires exposure to far fewer samples. By learning the mean target location for each underlying distribution, participants can use this knowledge to bias their estimates, particularly as sensory uncertainty increases. This model (‘Model 2’) predicts a smaller weight to the sensory information with increasing sensory uncertainty, and increased sensitivity to the underlying distributions as a function of exposure to the task. However, this model also predicts that for each level of sensory uncertainty, we should see identical weights being assigned to the sensory information for targets drawn from the two underlying distributions – the relative reliability of the underlying distributions is ignored by this model. This model is therefore still sub-optimal: the less variable (“narrow”) underlying distribution is a more reliable indicator of the likely target location across trials, and an optimal observer should take this into account.

Finally, a statistically efficient strategy would involve participants learning both the means and the relative variances of the underlying Gaussian distributions, and integrating this learned knowledge with the sensory information available on each trial in a Bayes-optimal manner (see Fig. 1B for an illustration). This model (‘Model 3’) predicts a drop in the weight to the sensory information with increasing sensory uncertainty, and a greater drop for trials in which the target is drawn from the narrow prior than when the target is drawn from the more variable (“broad”) prior distribution. Furthermore, as participants gain greater exposure to the underlying distributions, they should show a greater sensitivity to the statistical properties (means and relative variances) of the underlying distributions. Formally, if the mean and the variance of the sensory information on a given trial is given by *μ*_*l*_ and 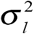, and the mean and the variance of the underlying trial-specific target distribution is given by *μ*_*p*_ and 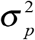, then the target location 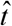 predicted by this model would be:

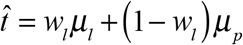

where *w*_*l*_, the weight assigned by the observer to the sensory information, should be:

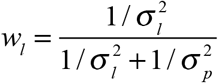

Previous research (Bejjanki et al., 2016) has shown that when adult participants are presented with this task, they behave in a manner that is consistent with the predictions of Bayes-optimal behavior (i.e., Model 3). Specifically, adults assigned a smaller weight to the sensory information as its reliability went down, and this drop was greater for trials in which the target is drawn from the narrow versus the broad prior distribution (Fig. S2). Furthermore, although participants learned the mean locations of the underlying distributions very rapidly, their weights continued to change throughout the experiment, suggesting that they learned the relative variances of the underlying distributions much more slowly.

#### Young children demonstrate sub-optimal behavior in this task

We examined the behavior of 6-7-year old children as they carried out 1600 trials of this task. We found that children in this age group behaved in a manner that was dramatically different from that observed previously with adults. Specifically, in both prior conditions (narrow and broad) and across all four temporal bins, children’s behavior was consistent with a strategy in which they based their estimates solely on the means of the sensory information available on each trial. They consistently assigned a weight near 1.0 to the centroid of the sensory information, and we found no reliable differences in the weights assigned to the sensory information irrespective of sensory uncertainty or prior reliability (Fig. 2A). In a 4 (temporal bin) x 3 (sensory uncertainty) repeated measures ANOVA of participants’ weights for trials in which the target was drawn from the broad prior distribution, there was no significant effect of temporal bin or sensory uncertainty, and no significant interaction (all *p*s > 0.05). Similarly, in a 4 (temporal bin) x 3 (sensory uncertainty) repeated measures ANOVA of participants’ weights for trials in which the target was drawn from the narrow prior distribution, there was no significant effect of temporal bin or sensory uncertainty, and no significant interaction (all *p*s > 0.1). Finally, in a 4 (temporal bin) x 2 (prior reliability) repeated measures ANOVA of weights in the high sensory uncertainty condition, we again found no significant effect of temporal bin or prior reliability, and no significant interaction (all *p*s > 0.8), further underscoring that children were not taking the reliability of the trial-specific prior distribution into account. This overall pattern of behavior is consistent with the predictions of Model 1 described above, and inconsistent with the predictions of Models 2 and 3 – 6-7-year-old children behave in a sub-optimal manner that ignores the uncertainty inherent in the two available sources of information (i.e., the likelihoods and the priors).

**Figure 2:**
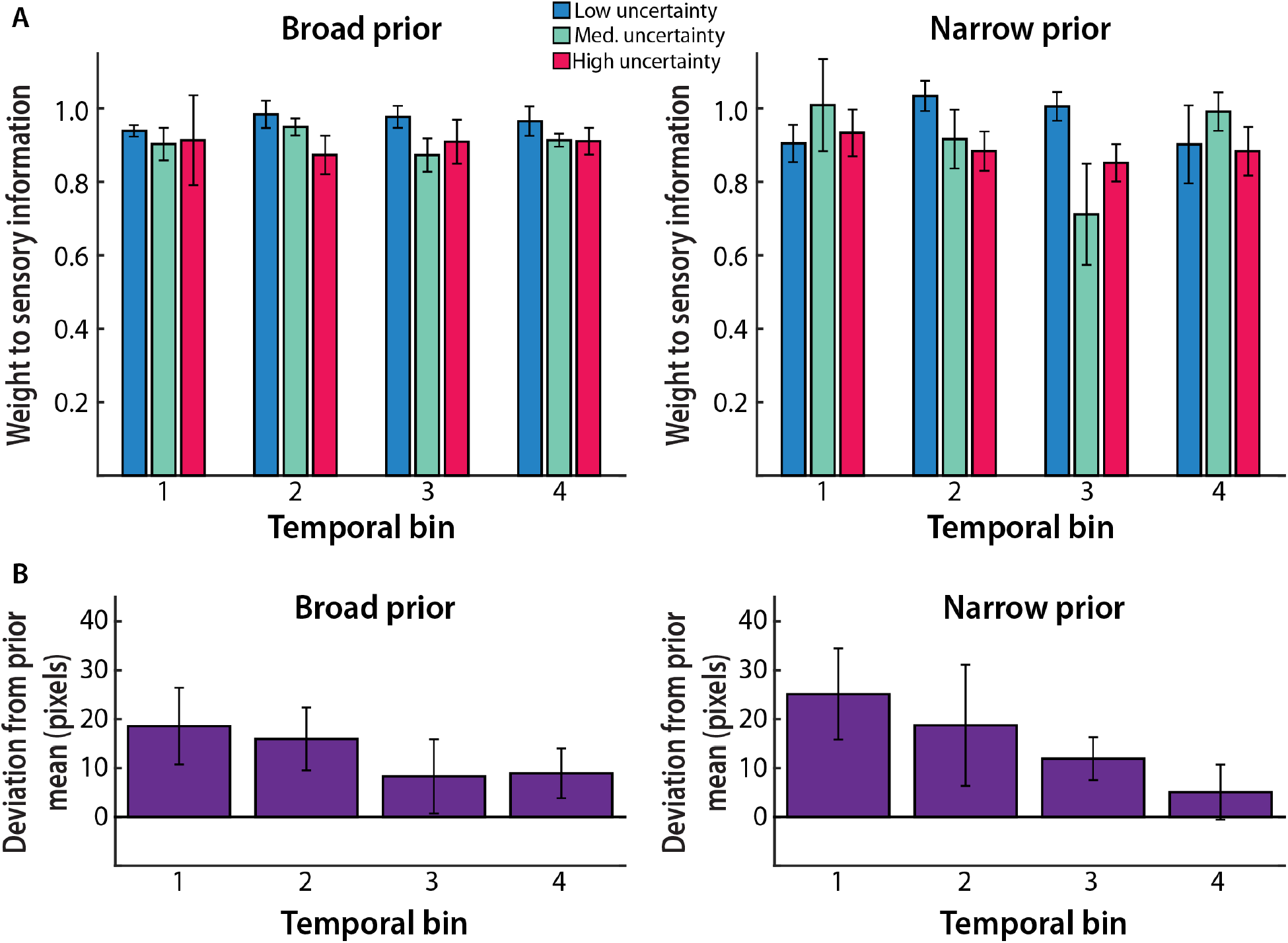
Learning and inference in Experiment 1. (A) Weights assigned to the centroid of sensory information. 6-7-year olds behaved in a manner dramatically different from that observed previously with adults (see Fig. S2), and inconsistent with the predictions of Bayes-optimal behavior. Each temporal bin includes 300 trials split between the two prior conditions and the four bins are depicted in temporal order. (B) Mean response locations in the absence of sensory information. Participants rapidly learned the true prior means for both the broad (left) and narrow (right) priors. For illustrative purposes, the y-axis represents deviation from the true mean, for each prior condition. Each temporal bin includes 100 trials split between the two prior conditions and the four bins are depicted in temporal order. Columns represent means and error bars represent SEM across participants.

We next examined the dynamics of learning in this task. Specifically, we considered the possibility that children might be learning the statistical properties of the underlying Gaussian distributions, despite being unable to appropriately use this information when computing weights (much as they were unable to appropriately take the sensory uncertainty available on each trial into account). We examined children’s behavior in the prior-only condition, in which they had to estimate the target location in the absence of any sensory information. We found that children were able to learn the means of the underlying distributions (Fig. 2B). In the first temporal bin (i.e., within the first 400 trials) children’s estimates of the target location were on average marginally different from the true prior mean locations (plus or minus noise estimated from the low likelihood variance conditions; broad prior: *t*_7_ = 1.94, *p* = 0.09; narrow prior: *t*_7_ = 2.26, *p* = 0.06), suggesting that they were yet to perfectly learn the mean locations. However, in all subsequent bins, their estimates of the target location were on average indistinguishable from the true prior mean locations (broad prior: all *p*s > 0.11; narrow prior: all *p*s > 0.12). This finding demonstrates that children in this age group are sensitive to, and capable of learning, at least one statistical property of the underlying distributions. It is intriguing that they nevertheless do not show any influence of this learned knowledge when weighting the two sources of information (for instance, in line with the predictions of Models 2 and 3).

## Experiment 2

In Experiment 1, we found that 6-7-year-old children, unlike adults, do not take sensory uncertainty into account despite it being available on each trial. Similarly, they do not take the variances of the underlying prior distributions into account despite learning at least one aspect (i.e., the mean) of these distributions. This pattern of behavior potentially provides further evidence for sub-optimality in integrating multiple sources of information at this age. However, there is an alternative to the hypothesis that young children are incapable of efficiently combining two or more sources of information: namely, the complexity involved in this task may have prevented them from utilizing an otherwise intact computational system for integrating uncertain sensory information with learned prior information. Specifically, in order to effectively track, learn and use the underlying distributions in this task, participants needed to be able to deploy sophisticated mechanisms of cognitive control and executive function (e.g., working memory, inhibitory control and cognitive flexibility). Prior research suggests that these mechanisms are still developing in children of this age group (Davidson, Amso, Anderson, & Diamond, 2006; Luciana & Nelson, 1998; Zelazo et al., 2013) (see Best and Miller (2010) and Carlson, Zelazo, and Faja (2013) for reviews), leading to the possibility that a deficit in these mechanisms may represent a key factor limiting young children’s behavior in this task.

To explore this possibility, we ran a follow-up experiment with a new group of 6-7-year-old children. We again used the spatial localization task – the goal was to estimate the location of a ‘hidden’ target based on uncertain sensory information presented in the form of a variable cluster of dots. However, a crucial difference was that we reduced task complexity by drawing targets from only one underlying Gaussian distribution – the narrow prior distribution used in Experiment 1 – located either in the left or the right half of the screen (counterbalanced across participants). All other aspects of the experiment, including the total number of trials, were identical to those in Experiment 1 (participants were thus exposed to twice as many samples from the single underlying distribution). Prior research (Bejjanki et al., 2016) has shown that when adult participants are presented with this simplified task, they continue to behave in a manner that is consistent with the predictions of Bayes-optimal behavior (Fig. S3). Adults not only assigned a lower weight to the sensory information as sensory uncertainty increased, but this drop was greater when targets were drawn from the more reliable underlying distribution.

Furthermore, participants rapidly learned the means of the underlying distributions, but learned the variances more slowly.

### Method

#### Participants

Eight 6- to 7-year old children (2 male; 6 female) participated in this experiment (none of whom participated in Experiment 1). Participants’ ages ranged from 6 years and 6 days to 7 years and 180 days (M = 6.92 years). Two additional participants were eliminated from the study due to an unwillingness to complete the task. Participants provided informed assent, and their caregivers provided informed written consent. Participants were given small prizes in the form of stickers, small toys etc., during their participation, while caregivers were compensated $20 per hour, for their time. Hamilton College’s institutional review board approved all experimental procedures.

#### Task description

The task procedure was identical to that used in Experiment 1, with the exception that the horizontal and vertical co-ordinates of the ‘hidden’ target location on each trial were independently sampled from a unimodal 2-D isotropic Gaussian distribution (rather than the bimodal distribution used in Experiment 1). The single underlying Gaussian distribution was centered in the left half of the display for half the participants, and in the right half for the other participants (with the locations being identical to the locations of the two distributions in Experiment 1). The variance of the distribution was identical to the variance of the less variable (“narrow”) underlying distribution used in Experiment 1 (i.e., it had an SD of 20 pixels).

#### Task procedure

Before the start of the experiment, participants were provided task instructions that were nearly identical to those in Experiment 1. The one exception was that participants were no longer told that the bucket could be located on either the left or the right side of the display, but rather that it would be located on one side or the other. On each trial, the procedure was identical to that in Experiment 1. Each participant again completed 1600 trials across two one-hour sessions – each session involved 800 trials and was run on a different day.

#### Data Analysis

As in Experiment 1, for the trials in which sensory information was available, we used linear regression to compute the weight (*w*_*l*_) assigned to the centroid of the dot cluster (the likelihood), with the weights assigned to the mean of the underlying distribution (the prior) being defined as (1-*w*_*l*_). For the trials in which no sensory information was available, we computed participants’ mean responses across all trials in each temporal bin. As in Experiment 1, we again focused on performance in the vertical dimension. Since all 1600 trials in this experiment involved targets being drawn from a single underlying distribution, participants were exposed to twice as many samples from the underlying distribution, compared to Experiment 1. Accordingly, the total number of trials were split into eight temporal bins (so as to include the same number of trials in each temporal bin as in Experiment 1).

### Results

#### Young children are capable of Bayes-optimal behavior when task complexity is reduced

If 6- to 7-year-old children are generally incapable of combining uncertain sensory information with learned prior information in a Bayes-optimal fashion, then we should continue to see sub-optimal behavior (as in Experiment 1) in this simplified task. Instead, we found that children demonstrated a pattern of behavior consistent with the predictions of Bayes-optimal behavior. Specifically, there was a reliable interaction between exposure to the task and sensory uncertainty: as participants gained more exposure to the task, they assigned a smaller weight to the sensory information, and this drop in weight was greater as the sensory information increased in uncertainty (Fig. 3A). In an 8 (temporal bin) x 3 (sensory uncertainty) repeated measures ANOVA of participants’ weights, there was a main effect of temporal bin, F (7,49) = 2.19, *p* = 0.05, a main effect of likelihood, F (2,14) = 60.86, *p* < 0.0001, and an interaction between the two factors, F (14,98) = 1.93, *p* = 0.03. Indeed, by the end of exposure to the task in Experiment 2 (i.e., in the final temporal bin), children assigned a reliably smaller weight to the sensory information as the uncertainty of sensory information increased across the three levels (1×3 repeated measures ANOVA: F (2,14) = 34.97, *p* < 0.0001; Fig. 4). Notably, the difference in behavior observed in Experiment 2, in comparison to Experiment 1, cannot be explained merely by the fact that participants were exposed to twice as many samples from the underlying distribution in Experiment 2. Even when they were exposed to an identical number of samples from the underlying distribution as in Experiment 1 (i.e., in the fourth temporal bin), children assigned a reliably smaller weight to the sensory information as the uncertainty of sensory information increased across the three levels (1×3 repeated measures ANOVA: F (2,14) = 17.42, *p* < 0.001). Furthermore, the children’ behavior was statistically indistinguishable from adult behavior in this task – comparing their behavior with adult behavior (using data from Bejjanki et al. (2016)), via a 2 (group: 6-7-year-old children, adults) x 8 (temporal bin) x 3 (sensory uncertainty) mixed ANOVA of participants’ weights revealed no significant effect of participant group, nor any interaction between group and any other factor (all *p*s > 0.34).

**Figure 3:**
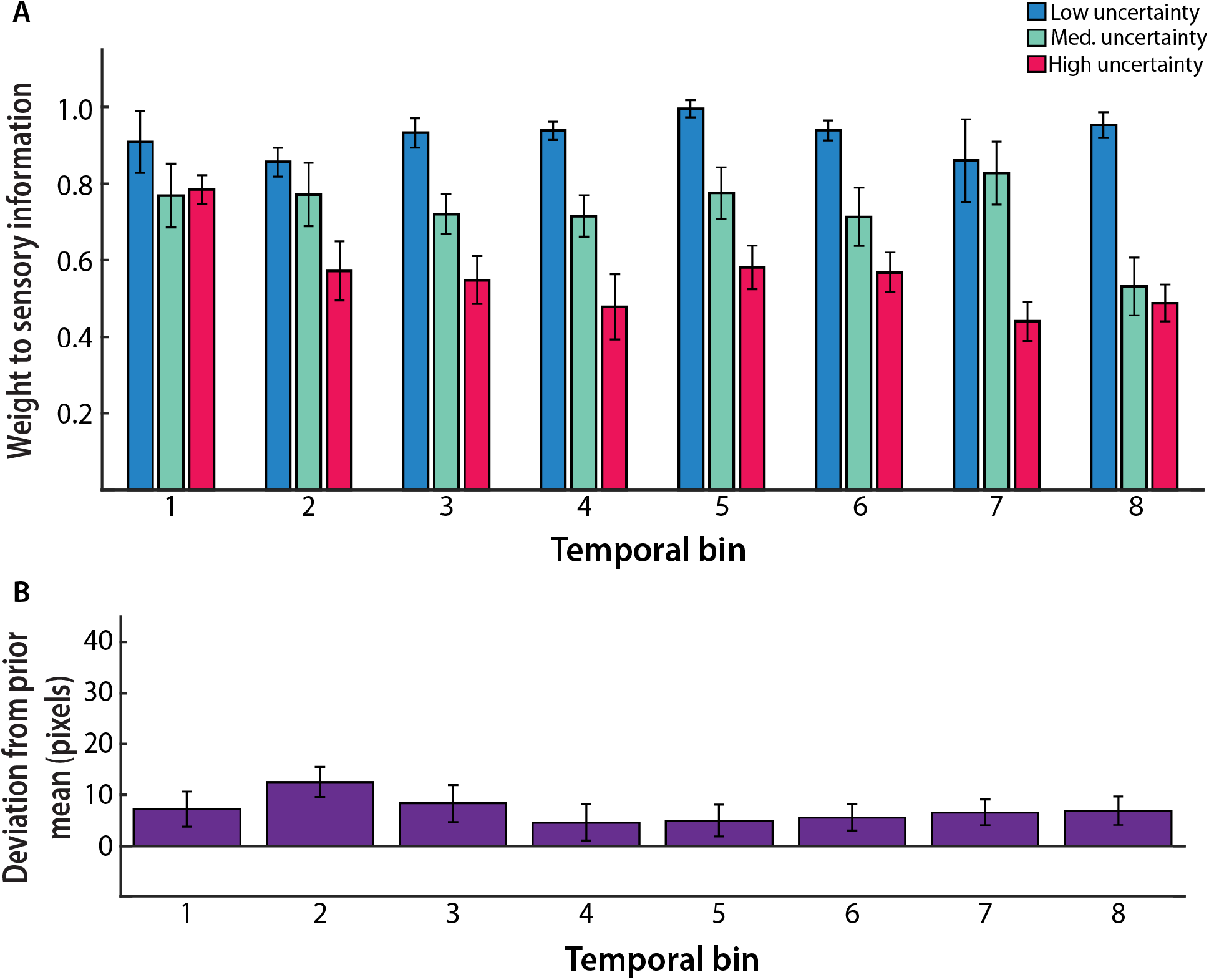
Learning and inference in Experiment 2. (A) Weights assigned to the centroid of the sensory information. Like adults (see Fig. S3) and consistent with the predictions of Bayes-optimal behavior, 6-7-year olds relied less on the sensory information (i.e., assigned a smaller weight to the centroid of the sensory information) as sensory uncertainty increased. Furthermore, this drop was greater as participants gained more exposure to the task. Each temporal bin includes 150 trials and the eight bins are depicted in temporal order. (B) Mean response locations in the absence of sensory information. Participants rapidly learned the true prior mean. For illustrative purposes, the y-axis represents the mean deviation from the true mean. Each temporal bin included 50 trials and the eight bins are depicted in temporal order. Columns represent means and error bars represent SEM across participants.

**Figure 4:**
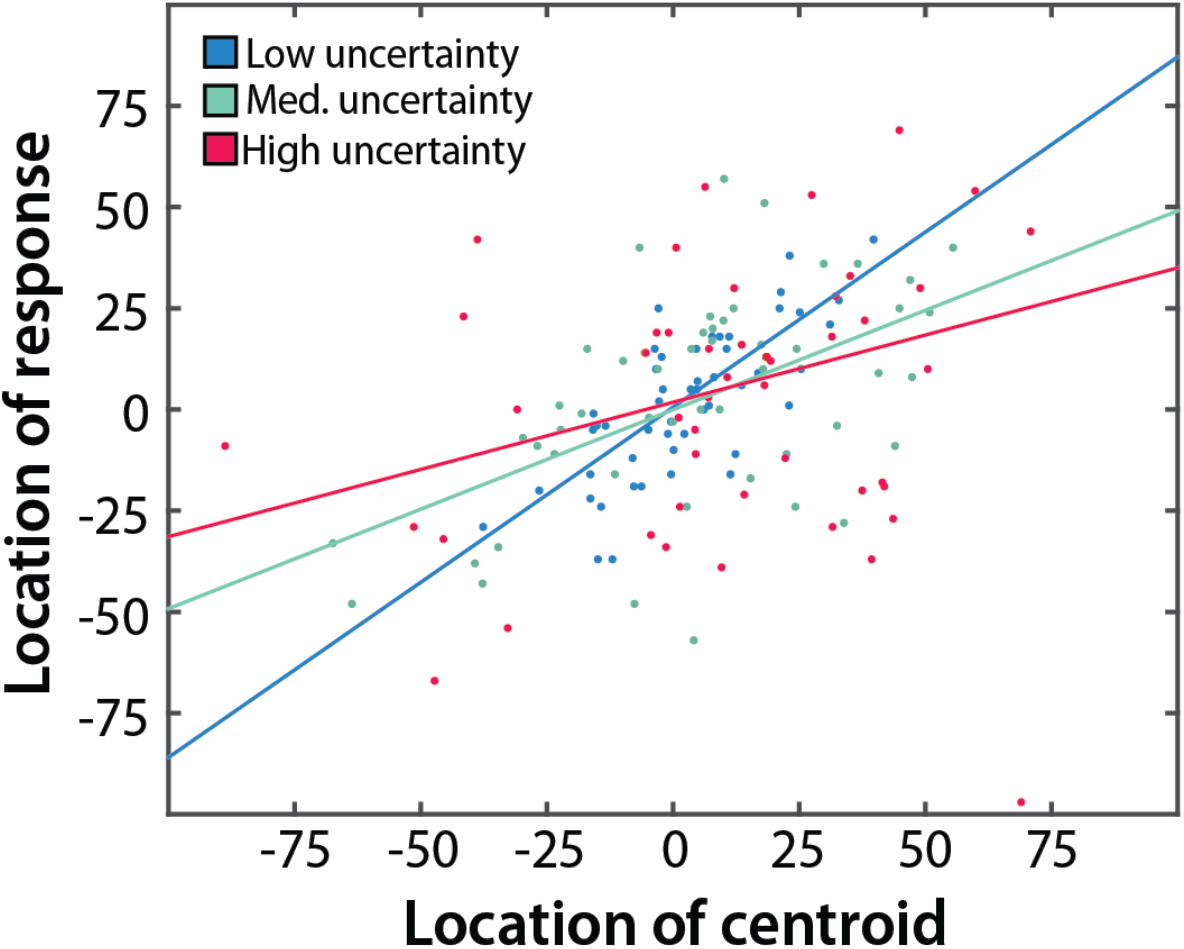
Data from the final temporal bin, for a representative participant in Experiment 2. Consistent with the predictions of Bayes-optimal behavior, as sensory uncertainty increased, participants shifted from selecting the centroid of the sensory information (the diagonal), towards selecting the mean of the underlying prior distribution (zero on the y-axis) as their estimate for the target location. For illustrative purposes, the mean of the underlying prior distribution was subtracted from both the response locations, and the centroid locations. Each dot represents a trial and solid lines represent the best-fit regression lines in each condition.

Similarly, in the prior-only condition, we found that children learned the true prior mean very rapidly (Fig. 3B) – within the first temporal bin, children’s estimate of the target location was on average indistinguishable from the true prior mean location (plus or minus noise estimated from the low likelihood variance conditions; *t*_7_ = 0.99, *p* = 0.35). Furthermore, in all subsequent bins except the second bin, children’s estimate of the target location continued to be indistinguishable on average from the true prior mean location (Bin 2: *t*_7_ = 3.55, *p* <0.01; Bins 3-8: all *p*s > 0.12). Furthermore, comparing their behavior with adult behavior (using data from Bejjanki et al. (2016)) via a 2 (group: 6-7-year-old children, adults) x 8 (temporal bin) mixed ANOVA revealed no significant effect of participant group, nor an interaction between group and temporal bin (all *p*s > 0.64), suggesting that children learned the true prior means at a rate that was on par with adults. Taken together, these findings suggest that children in this age group, like adults, have the ability to behave efficiently by taking the reliability of the likelihood (i.e., the variance of the sensory information) into account, and most importantly, by learning and integrating prior information with uncertain sensory information in a Bayes-optimal fashion.

## Discussion

Across two experiments, we used a spatial localization task to examine the extent to which 6-7-year old children are capable of efficiently combining uncertain sensory information available on each trial with prior information that could be learned over repeated exposure to the task. We demonstrate that children in this age group, like adults, are capable of carrying out statistically efficient computations. Specifically, we show that they are sensitive to the statistical properties of the task (i.e., the reliability of sensory and prior information), and they are able to track and learn these statistics at a rate on-par with that of adults. Importantly, we also show that children in this age group are able to integrate learned prior information with uncertain sensory information on a trial-by-trial basis, and in a manner consistent with the predictions of Bayes-optimal behavior. Prior research has shown that young children and infants are sensitive to the statistical properties of stimuli in their environment (Ernst & Banks, 2002; Gopnik et al., 2001; Jusczyk & Aslin, 1995; Kirkham et al., 2002; Kuhl & Meltzoff, 1982; Neil et al., 2006; Saffran et al., 1996; Smith & Yu, 2008; Xu & Garcia, 2008), and the ability to appropriately learn and use such environmental regularities is involved in the development of a number of cognitive abilities (e.g., object recognition, language acquisition, causal inference etc.). Our findings extend this prior research by further showing that young children are capable of utilizing such learned regularities in a statistically optimal fashion.

However, we also show that in a more complex version of the same task, children at this age behave sub-optimally, and in a manner dramatically different from adult behavior. Specifically, when faced with uncertain sensory information regarding stimuli drawn from a mixture of two underlying prior distributions, children, but not adults, consistently chose the centroid of the sensory information as their estimate for the target location, irrespective of sensory uncertainty or prior reliability. Notably, children exhibited this behavior despite learning the mean locations of the underlying distributions, and having access to the reliability of the sensory information on each trial. This pattern of behavior is consistent with an inability to appropriately take each available source of information into account, which could be due to deficits in the computation of task-relevant variables (including likelihoods, priors, objective functions and decision rules), or limitations in the neural implementation or approximation of Bayesian inference (Drugowitsch, Wyart, Devauchelle, & Koechlin, 2016; Rahnev & Denison, 2018). Note that one can only combine information across multiple cues if one can keep track of which aspects of that information should be combined and under what conditions. For example, in Experiment 1, participants were required to simultaneously encode three levels of sensory uncertainty, the presence of two priors, and that each prior was reliable to a different extent. In contrast, in Experiment 2, while the same three levels of sensory uncertainty were present, there was only one prior with one level of uncertainty. Thus, the results from our two experiments suggest that while young children are capable of carrying out Bayes-optimal computations in simplified situations, this ability is severely influenced by task complexity, and when overwhelmed by that complexity, children revert to a simpler implicit strategy for guiding their performance (i.e., selecting the centroid of the sensory information irrespective of sensory or prior uncertainty).

Our findings have important implications for how we should interpret failures in young children’s ability to carry out sophisticated computations. For instance, in a recent study, Chambers et al. (2018) used a similar spatial localization task to examine the extent to which 6-11-year-olds and adults combined uncertain sensory information with prior information. Notably, targets were drawn from one of two underlying Gaussian distributions in a *blocked* fashion (via four alternating blocks of 120 trials), and they did not examine learning (a cue to the block-relevant prior was always visible on the screen). They found that 6-8-year-old children, like adults, appropriately assigned a smaller weight to the sensory information as sensory uncertainty increased, but unlike adults, they sub-optimally did not take the relative reliability of the block-specific underlying distribution into account. The authors concluded that Bayes-optimal behavior may not be inherent but must at least partly be learned during development. Our findings suggest an alternative interpretation: rather than being incapable of Bayes-optimal computations, young children might have responded to the complexity of their task by reverting to a computationally simpler but sub-optimal strategy, with task complexity affecting young children’s behavior to a greater extent than that of adults. Indeed, the Chambers et al. (2018) study involved a task with an intermediate level of complexity (compared to the two experiments in our study), so it is interesting to note that 6- to 8-year-olds demonstrated a pattern of behavior that was intermediate in sophistication (consistent with predictions of Model 2 described above), compared to that observed in the two experiments in our study.

How might task complexity differentially influence young children’s ability to carry out Bayes-optimal computation? One possibility is that performance in tasks involving greater complexity (such as that seen in Experiment 1) necessitates the deployment of sophisticated mechanisms of cognitive control and executive function. Specifically, in order to effectively track, learn and use the underlying distributions, participants needed to be able to hold in memory, mentally manipulate, and act on the basis of both their evolving knowledge of the prior and the sensory information available on each trial. In addition, on each trial, the hidden target was drawn from one of two underlying distributions, characterized by different means and variances, requiring participants to learn and maintain multiple “rules” in their mind, and to quickly and flexibly adapt their behavior to the changing situation (e.g., via inhibition and cognitive flexibility). There is substantial evidence that such top-down mechanisms show a protracted developmental progression, with performance in some tasks not reaching adult-like levels even by the teen years (Best & Miller, 2010; Carlson et al., 2013; Davidson et al., 2006; Luciana & Nelson, 1998; Zelazo et al., 2013). The inability to utilize these mechanisms early in development therefore represents an important limitation that young children, but not adults, may be subject to.

A large body of previous research has established that human behavior in a range of tasks might look sub-optimal (or “irrational”) if the cognitive and computational limitations that observers are subject to are not appropriately accounted for (Lieder & Griffiths, 2019; Simon, 1956; Tversky & Kahneman, 1974). In the current context, several studies have shown that young children’s behavior in tasks drawing upon top-down mechanisms, such as cognitive control and executive function, is critically moderated by task complexity. For instance, Luciana and Nelson (1998) found that while 5-7-year olds were indistinguishable from adults when carrying out simple versions of a spatial working memory task, as task demands became more rigorous, such as when they had to simultaneously recruit mnemonic traces, sequence motor responses and organize behavior using a strategy or hierarchical series of goals, performance in 5-7-year olds, but not adults, deteriorated rapidly. Similarly, Davidson et al. (2006) found that while even 4-year olds could simultaneously hold information in mind and inhibit a dominant response when rules remained constant, the ability to flexibly switch between rules was not adult-like even in 13-year-olds. The sub-optimal behavior observed in Experiment 1 may thus be less about an inability to carry out Bayes-optimal computations (i.e., weighting each available cue by its relative reliability), and more about a deficiency in the top-down skills needed to effectively deploy this ability. In this scenario, the “failures” in statistically efficient computations observed in young children need not necessarily be attributed to deficits in the fundamental computational capacity available to children early in development, but rather to ancillary immaturities in general cognitive abilities that mask the operation of these sophisticated computations.

Future work might explore the extent to which our findings are applicable to other tasks involving the integration of sensory information with learned prior information. For instance, an open question is whether behavior in all tasks that involve flexibly utilizing multiple learned prior distributions involves the deployment of top-down control mechanisms. A large body of previous research has established a distinction between controlled and automatic processing (Shiffrin & Schneider, 1977), with controlled processing involving pre-frontally mediated top-down control (Miller & Cohen, 2001). As such, one possibility is that flexibly utilizing multiple prior distributions that are well-learned (in contrast to the novel distributions learned in the current task) might represent an instance of automatic processing that precludes the deployment of top-down control mechanisms, and would therefore not be subject to the influence of complexity as demonstrated here.

Finally, it is important to note that the current study speaks most directly to the specific problem of efficiently integrating learned prior information with uncertain sensory information. Most previous studies reporting that young children differ substantially from adults in their ability to utilize multiple sources of information have instead considered tasks involving simultaneously available cues (e.g., from touch and vision). Our findings cannot speak to why young children fail to exhibit Bayes-optimal behavior in such tasks when combining two simultaneously available sensory cues. For instance, previous research has suggested that different neural regions, and patterns of interaction among them, may be involved in the integration of information in these two scenarios – frontal areas of the brain, such as the PFC and the lateral OFC, when combining prior information with sensory information (Nogueira et al., 2017; Vilares et al., 2012), versus low-level sensory areas when combining simultaneously available cues (Ban, Preston, Meeson, & Welchman, 2012; Gu, Angelaki, & DeAngelis, 2008). Furthermore, given that all the relevant information is available on each trial, experiments involving the combination of simultaneously available cues have minimal executive function and memory/learning demands. As such, the question of task complexity seems much less relevant than it is in our experiments. Conversely, mechanisms (e.g., cue recalibration or prioritizing the earliest available cue) implicated in the sub-optimal behavior observed when integrating multiple simultaneously available cue are less likely to be impediments to efficiently integrating sensory and prior information in a single modality and in the same frame of reference, as in our experiments. Similarly, when faced with multiple simultaneous cues, a key problem is inferring the extent to which both cues pertain to the same underlying stimulus (or source) – only cues pertaining to the same source should be integrated. This causal inference problem is much less relevant in situations involving the integration of prior and sensory information, since it is clear that the two cues pertain to the same underlying stimulus variable (in our case, the location on the touch screen). Teasing apart the similarities and differences between tasks involving two simultaneous sources of information and tasks involving sensory and prior information remains an important goal for understanding the underlying computational mechanisms, and their constraints, during development.

## Acknowledgements

This work was supported by a research grant from NIH (HD-037082; Richard Aslin, PI). We are grateful to Holly Palmeri and Emily Williams for their help with recruiting and running participants.

## Author Contributions

V.R.B. and R.N.A. conceived the study; V.R.B. and E.R.R. conducted the experiments and analyzed the data; V.R.B. and R.N.A. wrote the paper with assistance from E.R.R.

## Data Availability Statement

The data that support the findings of this study are available from the corresponding author upon reasonable request.

