## Supplementary Information for "Young children combine sensory cues with learned information in a statistically efficient manner: But task complexity matters"

### Table of contents

Supplementary Figure S1

Supplementary Figure S2

Supplementary Figure S3

**Supplementary Figure S1**

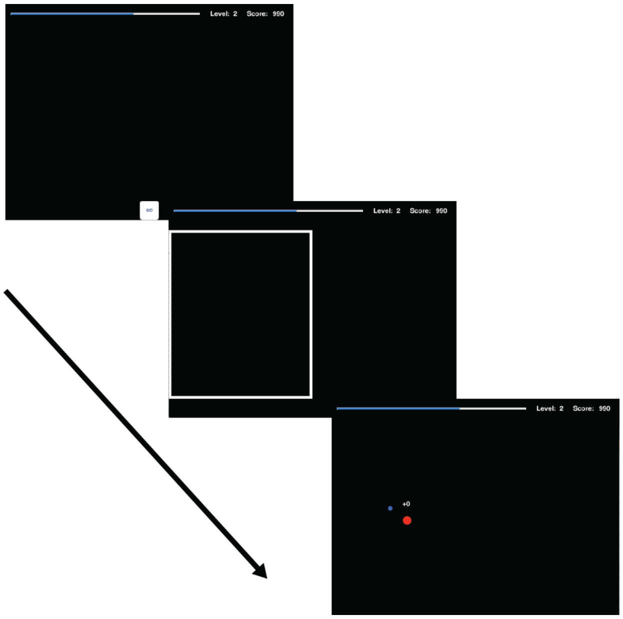


**Figure S1: The ‘prior-only’ condition.** An illustration of a typical trial. In this condition, participants localized the hidden target in the absence of any sensory information. After they touched a ‘Go’ button, participants saw a briefly flashed rectangle on either the left or right half of the screen (rectangle shown here on the left half) – they were told that the target was located within the rectangle. Since the two underlying distributions were centered at different locations on the screen (one in the left half and the other in the right half), the rectangle indicated which of the two underlying distributions the target on the current trial was drawn from. To further signal the relevant underlying distribution, the rectangle was either white or green in color. Feedback was provided post-touch.

**Supplementary Figure S2**

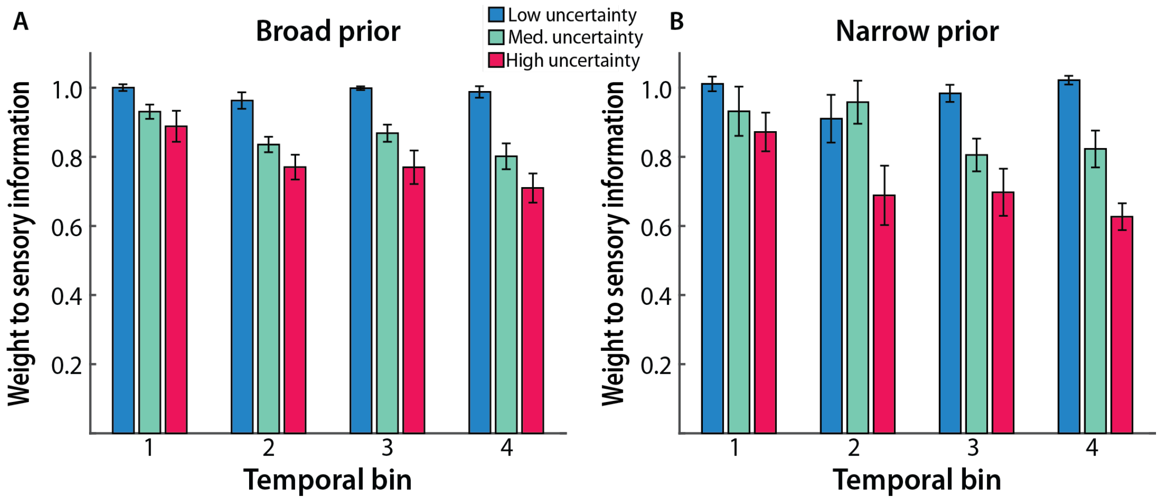


**Figure S2: Weights assigned by adult participants in the task used in Experiment 1.** In this task, consistent with the predictions of Bayes-optimal behavior, adult participants relied less on the sensory information (i.e., assigned a smaller weight to the centroid of the sensory information) as sensory uncertainty increased. This was true for both broad (A) and narrow (B) prior conditions, with the drop in the weight to the sensory information being greater for the narrow versus the broad prior condition. Furthermore, for both prior conditions, the drop in the weight to the sensory information as a function of increasing uncertainty, was greater as participants gained more exposure to the task. Each temporal bin includes 300 trials split between the two prior conditions and the four bins are depicted in temporal order. Columns represent means and error bars represent SEM across participants. Figure adapted from Bejjanki et al. (2016).

**Supplementary Figure S3**

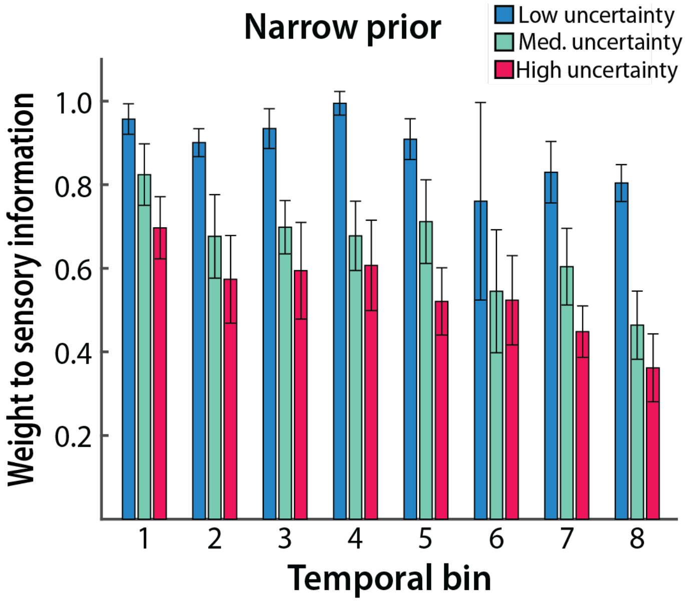


**Figure S3: Weights assigned by adult participants in the task used in Experiment 2.** In this simpler task, again consistent with the predictions of Bayes-optimal behavior, adult participants relied less on the sensory information (i.e., assigned a smaller weight to the centroid of the sensory information) as sensory uncertainty increased. Furthermore, the drop in the weight to the sensory information as a function of increasing uncertainty, was greater as participants gained more exposure to the task. Each temporal bin includes 150 trials and the eight bins of trials are depicted in temporal order. Columns represent means and error bars represent SEM across participants. Figure adapted from Bejjanki et al. (2016).
